# Integrating computational chemistry and machine learning to predict KRAS mutation-induced resistance

**DOI:** 10.64898/2026.04.10.717640

**Authors:** Katarzyna Mizgalska, Konstancja Urbaniak, Denis J. Imbody, Eric B. Haura, Wayne C. Guida, Sergio Branciamore, Aleksandra Karolak

**Affiliations:** Department of Machine Learning, Moffitt Cancer Center, Tampa, FL, United States of America; Department of Chemistry, University of South Florida, Tampa, FL, United States of America; Department of Computational and Quantitative Medicine, City of Hope, Monrovia, CA, United States of America; Department of Thoracic Oncology, Moffitt Cancer Center, Tampa, FL, United States of America

## Abstract

Mutation-induced drug resistance is a major contributor to the failure of targeted cancer therapies, particularly in tumors driven by mutations in the KRAS oncogene. Although covalent inhibitors effectively target KRAS G12C, secondary mutations such as G12C/Y96C, G12C/Y96S, and G12C/Y96D lead to resistance despite leaving the covalent attachment site intact. To predict these resistance outcomes, we developed a computational framework that integrates molecular dynamics-derived structural, energetic, thermodynamic, and contact-based descriptors with machine learning. Features extracted from simulations of treatment-sensitive and treatment-resistant KRAS mutants were used to train logistic regression, random forest, support vector machine, and Bayesian Network classifiers, achieving average accuracies above 90%. Solvent-accessible surface area variability, Lennard-Jones 1,4 energy, mean square displacement, and root mean square fluctuation emerged as the most discriminatory features. Residues G10, E62, and H95 showed the highest predictive value. This approach highlights conformational and solvent-exposure changes as central drivers of KRAS drug resistance and provides a generalizable workflow for other clinically relevant mutant targets.

**Author Summary:** Mutation-induced resistance is a common challenge across many cancer types and is often associated with aggressive tumor progression and poor therapeutic response. Investigating the dynamic properties of proteins harboring such mutations provides valuable insights into the structural and functional consequences of these alterations, thereby helping to elucidate the mechanisms of drug resistance. Machine learning algorithms are particularly effective at uncovering complex patterns within high-dimensional data, such as molecular dynamics simulation trajectories. Integrating these algorithms with analysis of protein dynamics holds significant potential to aid in drug discovery challenges by reducing both time and resource demands while increasing the likelihood of identifying effective therapeutic candidates. As a proof of concept, we developed a computational framework that integrates molecular dynamics-derived molecular features with machine learning to distinguish treatment-sensitive from treatment-resistant KRAS mutants. KRAS is known for drug resistance arising from secondary mutations that disrupt inhibitor binding despite intact covalent attachment sites. The models achieved over 90% accuracy and identified solvent-exposure and conformational changes at residues G10, E62, and H95 as key predictors of treatment resistance. This workflow offers a generalizable strategy for understanding and forecasting mutation-induced resistance.

## Introduction

Mutation-induced drug resistance presents a major challenge in cancer therapy, particularly for targeted treatments. Such resistance can arise through various mechanisms [1], including activation of alternative signaling pathways, eg, BRAF(V600E) inhibition by vemurafenib in colon cancer [2]; impaired apoptosis, as seen in p53(R273W) resistance to 5-fluorouracil [3]; and enhanced DNA repair, eg, BRCA1/2 reversion mutations conferring resistance to polyadenosine diphosphate ribose polymerase inhibitors [4]. This study focuses on a common mechanism of resistance driven by direct mutations in the drug target itself. Such mutations frequently alter the chemical and conformational properties of the target, often within the binding site, disrupting drug-receptor interactions. This mechanism, among others, is especially concerning for one of the most frequently mutated oncogenes, Kirsten rat sarcoma viral oncogene homolog (*KRAS*) [5, 6].

The *KRAS* oncogene encodes a 21 kDa guanosine triphosphate hydrolase (GTPase) K-Ras (KRAS henceforward) that regulates key signaling pathways involved in cell-cycle control [6, 7]. Structurally, KRAS consists of a G-domain (residues 1–166), which binds guanine nucleotides, and a hypervariable region (residues 167–189), which mediates membrane association [7]. Its secondary structure includes 6 β-sheets and 5 α-helices connected by flexible loops. Important regions (shown in **Fig 1**) include the switch I and switch II loops, which interact with effector and regulatory proteins, and the P-loop, which binds nucleotide phosphate groups [7]. KRAS homolog alternates between an inactive guanosine diphosphate– (GDP) bound and an active guanosine triphosphate– (GTP-) bound state. Guanine nucleotide exchange factors (GEFs) promote the release of GDP, allowing GTP, which is more abundant in the cell, to bind and stabilize the active conformation [8]. Binding of GTP activates KRAS by inducing conformational changes in the switch I and switch II loops. The activated protein then engages downstream signaling pathways, including the mitogen-activated protein kinase (MAPK) pathway, the Ras-like small GTPases pathway, and the phosphoinositide-3-kinase pathway, which are critical regulators of cell proliferation, survival, growth, and differentiation [9]. Termination of signaling requires GTP hydrolysis, facilitated by GTPase-activating proteins, returning KRAS to its inactive state [8].

**Fig 1.**
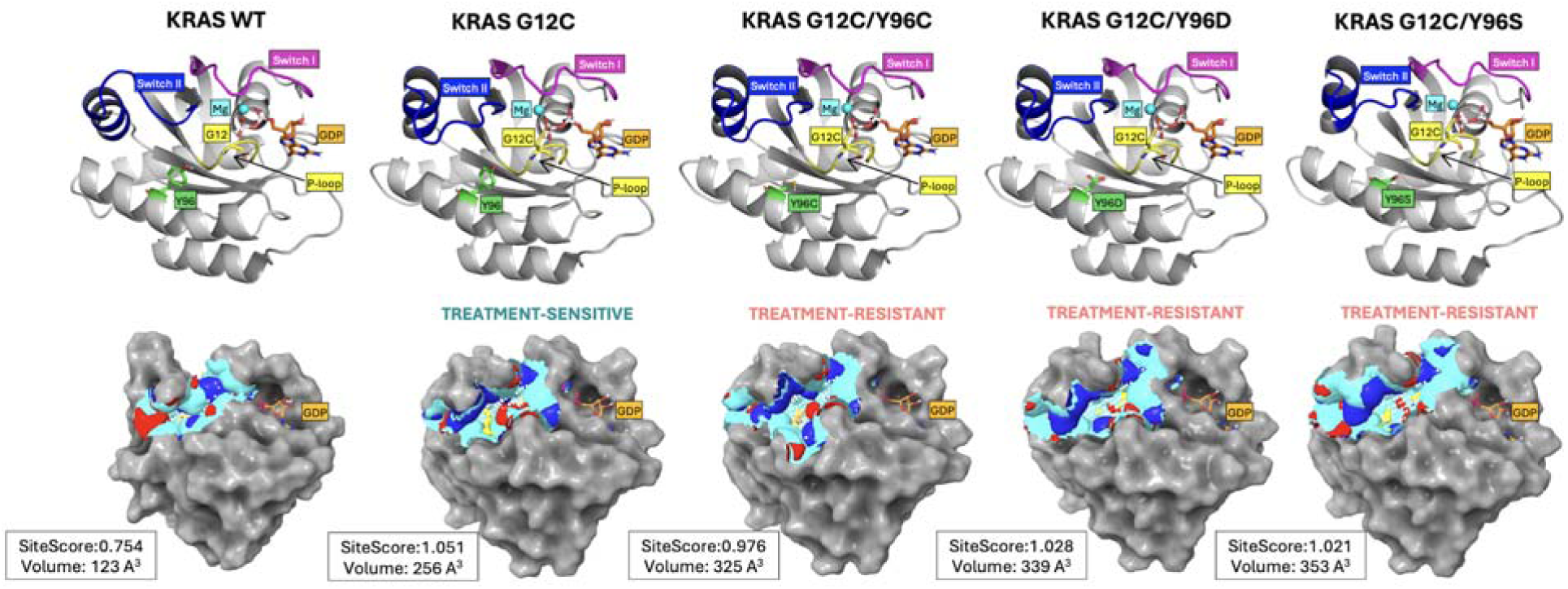
KRAS mutation points and SiteMap analysis. The first row shows labeled structural elements of KRAS forms; the second row shows their respective switch II binding sites. Binding site properties: binding site surface (cyan), hydrogen bond donors (blue), hydrogen bond acceptors (red), hydrophobic surface (yellow), and GDP (orange). The binding SiteScore and volume are reported next to each molecule. For visualization purposes, switch II is approximated on residues 58–72, switch I on residues 30–40, and P-loop on residues 10–14 as suggested in the following work [7]. The visualizations were prepared with PyMOL Molecular Graphics System [40]. Abbreviations: GDP, guanosine diphosphate; KRAS, Kirsten rat sarcoma viral oncogene homolog. Corresponding PDB structures include 6MBT (KRAS WT); 6OIM (KRAS G12C); 6OIM manually mutated (KRAS G12C/Y96C/D/S).

Mutations in signaling pathways such as Ras/Raf/MAPK are frequently implicated in cancer, with KRAS being one of the most commonly mutated oncogenes [6] accounting for 11.6% of all carcinomas [5]. In pancreatic ductal adenocarcinoma, KRAS mutations are present in 81.7% of cases, with G12D as the predominant subtype [5]. In colorectal cancer, KRAS mutations occur in 38% of patients, primarily G12D and G12V[5], while in non–small-cell lung cancer, the prevalence is 21.2%, with G12C representing 45% of cases [5, 6, 10, 11]. Point mutations at KRAS residues G12, G13, and Q61 impair GAP-mediated GTP hydrolysis, leading to constitutive KRAS activation and tumorigenesis [12].

Until recently, KRAS-driven tumors were considered difficult to treat because of KRAS’s picomolar affinity for GTP and the absence of other druggable pockets on its surface. As a result, therapeutic strategies focused on targeting KRAS indirectly [13, 14]. A breakthrough came in 2013, when Ostrem et al [15] reported the first covalent inhibitor targeting the KRAS G12C mutant. The compound bound specifically to a cysteine residue within an allosteric pocket located beneath the switch II region in the inactive (GDP-bound) KRAS conformation. This breakthrough led to the development of the first KRAS G12C inhibitors approved for clinical use. In 2021, AMG510 [16] (sotorasib, Lumakras) was approved by the FDA for treating non–small-cell lung cancer, followed by MRTX849 [17] (adagrasib, Krazati) in 2022. Both drugs irreversibly bind to the cysteine residue at position 12 and form noncovalent interactions with other amino acids in the switch II binding site, including tyrosine (Y96), glutamine (Q99), and histidine (H95). This interaction network stabilizes the inactive, GDP-bound conformation of KRAS, thereby inhibiting oncogenic signaling [18, 19].

Despite initial clinical success, resistance to sotorasib and adagrasib has emerged through various mechanisms [20, 21]. One involves RTK-mediated activation of wild-type RAS [22]. Another features secondary mutations in KRAS G12C, such as Y96D [23–25], Y96S [24, 25], and Y96C [24, 26], which confer resistance to both approved drugs. Other mutations, such as G12C/G13D, R68M, A59S, and A59T, are resistant to sotorasib alone, while Q99L is associated with resistance to adagrasib [24, 25]. These secondary mutations reactivate ERK signaling and are likely to promote broader resistance across tumor-cell populations [27].

Current efforts to target KRAS-driven tumors include inhibitors of GDP-bound KRAS G12C and G12D, as well as proteolysis-targeting chimeras for KRAS G12D and G12V, all in clinical trials [28]. Alternative strategies focus on the active, GTP-bound form of KRAS and binding regions outside of switch II to circumvent resistance from secondary mutations [29]. However, the molecular basis of resistance arising in the inactive-state binding site, particularly in KRAS secondary mutants, remains poorly understood, underscoring the potential benefit of targeted approaches against these specific variants.

This study moves beyond prevailing studies focused on conformational analyses [30], single KRAS mutants [31–33] or drug design [34]. Instead, we investigate the underlying factors of mutation-induced resistance caused by secondary mutations in the KRAS switch II binding site: KRAS G12C/Y96D, G12C/Y96C, and G12C/Y96S, which exhibit resistance to both sotorasib and adagrasib. As both drugs interact with residue Y96 [19], substitution at this site represents a direct structural change relative to the G12C single mutant, which may influence drug binding. However, other molecular features may also contribute to resistance. To address this, we developed a framework combining molecular dynamics– (MD-) derived descriptors with machine learning (ML) algorithms to identify features that differentiate treatment-resistant KRAS mutants from treatment-sensitive ones.

Integrating dynamic properties of biological systems with ML offers a novel approach to tackling complex challenges in drug discovery [35]. While some studies have incorporated MD descriptors into ML models, they have primarily focused on ligand-based pharmacophore analyses [36], without accounting for protein or binding site dynamics. Other approaches describe protein-binding sites statically and apply ML to support ligand design [37]. Although a few efforts have incorporated protein flexibility and ML, eg, to identify druggable binding sites [37–39], this approach has not yet been applied to address complex problems such as mutation-induced drug resistance.

## Results

This study investigated mutation-induced drug resistance using the KRAS oncogene as a case study. Based on experimental treatment responses, KRAS structures were classified either as treatment-sensitive (G12C) or treatment-resistant (G12C/Y96C, G12C/Y96S, and G12C/Y96D). Because the secondary mutations occur within the switch II binding site, we first assessed structural differences in this region using SiteMap (**Fig 1**). One key output, SiteScore, reflects the likelihood of small-molecule binding, with values near 1 indicating promising binding sites. KRAS WT exhibited a shallow switch II pocket with a low SiteScore (0.754) and small volume (123 Å³), suggesting unlikely small-molecule binding, consistent with the absence of endogenous ligands in native KRAS. In contrast, the G12C single mutant and all secondary mutants had a SiteScore near 1, but their binding pocket volumes varied: G12C had a smaller volume (256 Å³), while the secondary mutants ranged from 325 to 353 Å³. This volume increase is likely attributable to substitution of bulky tyrosine at position 96 with smaller residues (cysteine, serine, or aspartic acid).

To capture dynamic and subtle structural changes not evident from static analysis, we performed MD simulations of each KRAS mutant in the absence of inhibitors. From the MD trajectories, we extracted a comprehensive set of molecular descriptors reflecting global structural behavior and localized changes within key regions of the protein. These features were categorized into 4 groups: structural, energetic, thermodynamic, and intra/intermolecular contacts. The descriptors most strongly associated with resistance were related to the site of secondary mutation (residue 96), and the average solvent-accessible surface area (SASA) of residues G10 and A11. After removing highly intercorrelated variables (see Methods section), the remaining features were subjected to statistical and ML analyses.

### Univariate statistical analysis

The RMSD relative to the reference structure was calculated to assess system stability. Protein backbone RMSD plots (**Fig 2**) illustrate structural fluctuations in treatment-sensitive and treatment-resistant KRAS systems throughout the simulation. The treatment-sensitive systems reached a maximum RMSD of 0.41 nm (**Fig 2A**), while the treatment-resistant systems exhibited greater protein fluctuations, with RMSD values rising to 0.50 nm (**Fig 2B**). To better characterize this variability, we generated violin plots of backbone RMSD distributions for the most populated conformations (**Fig 2C**). In both groups, medians were broadly distributed around similar values (0.20 and 0.21 nm), although the treatment-resistant group exhibited greater variability than the treatment-sensitive group (RMSD range: 0.13–0.39 nm vs 0.13–0.32 nm). However, the Mann–Whitney U test revealed no statistically significant difference (*P* value >.05) in backbone RMSD distributions between groups.

**Fig 2.**
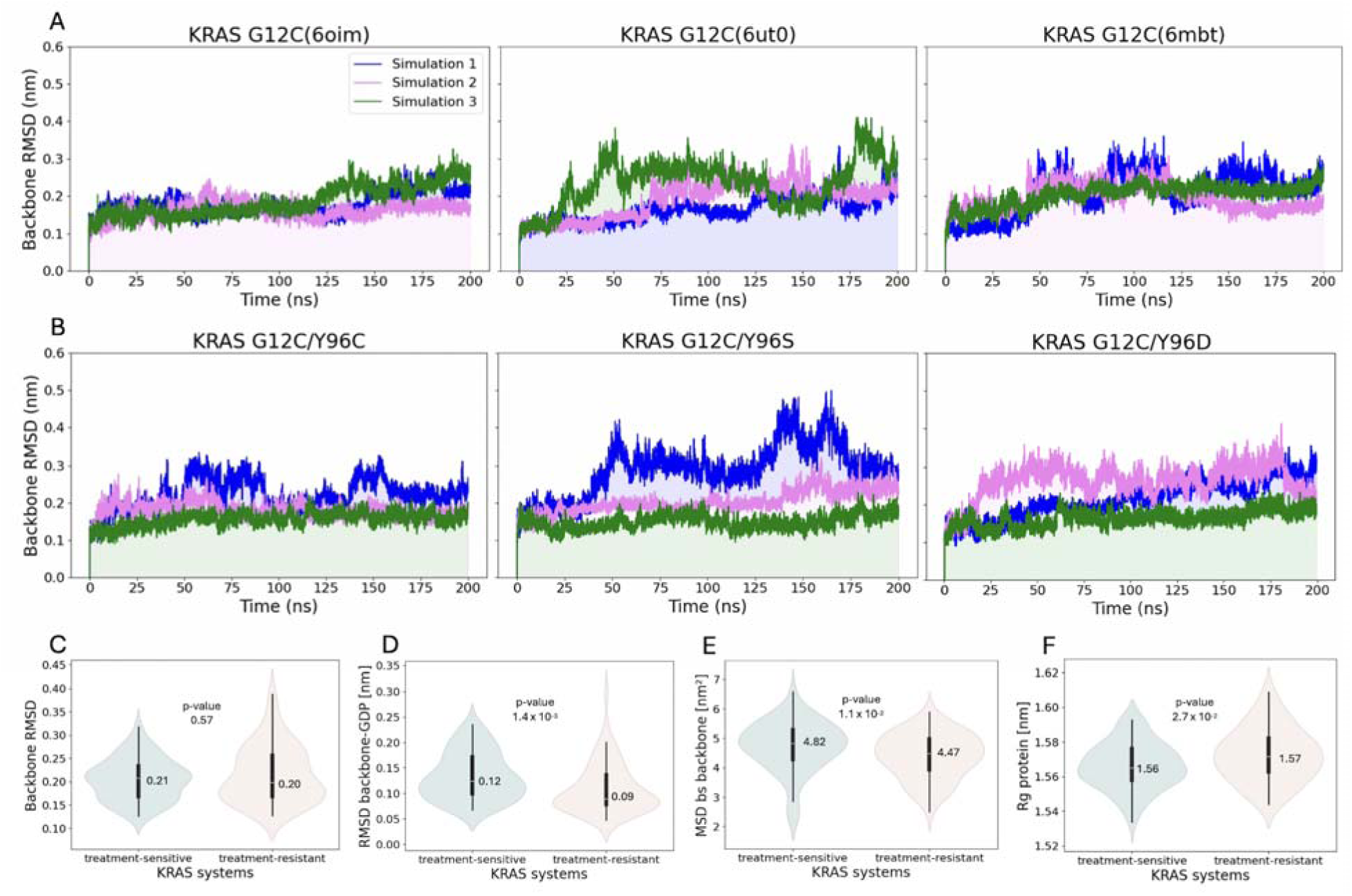
RMSD over the molecular dynamics simulation. (**A**) KRAS treatment-sensitive systems. (**B**) KRAS treatment-resistant systems. Data distribution of structural features with annotated median and P values. (**C**) Protein backbone RMSD. (**D**) Protein backbone–GDP RMSD. (**E**) Protein backbone MSD of switch II binding site residues. (**F**) Protein Rg. Abbreviations: GDP, guanosine diphosphate; KRAS, Kirsten rat sarcoma viral oncogene homolog; MSD, mean square deviation; RMSD, root mean square deviation.

Among all RMSD-based features, only the RMSD of the backbone–GDP complex showed a statistically significant difference (**Table 1**). While both groups displayed values concentrated near their medians (0.12 and 0.09 nm, **Fig 2D**), the treatment-sensitive systems were shifted toward higher RMSD values (mean: 0.13 vs 0.11 nm, mean values not shown) than treatment-resistant systems, suggesting their greater structural flexibility regarding the protein–GDP complex.

**Table 1.** The results of Mann–Whitney U test. . Molecular features with *P* value less than 0.05.

| Molecular feature | P-value |
| --- | --- |
| SASA G10 SD* | $1.50 \times 10^{-17}$ |
| SASA H95 AV* | $1.75 \times 10^{-12}$ |
| LJ 1,4 | $2.19 \times 10^{-8}$ |
| SASA E62 SD | $6.00 \times 10^{-8}$ |
| RMSF M72 | $1.20 \times 10^{-7}$ |
| SASA I100 AV | $1.35 \times 10^{-7}$ |
| SASA V103 SD | $1.46 \times 10^{-7}$ |
| SASA V9 AV | $1.83 \times 10^{-7}$ |
| SASA Q99 AV | $4.72 \times 10^{-6}$ |
| SASA K16 AV | $1.44 \times 10^{-5}$ |
| RMSF Q99 | $1.64 \times 10^{-5}$ |
| RMSF H95 | $2.83 \times 10^{-5}$ |
| RMSF Y64 | $4.62 \times 10^{-5}$ |
| Hb protein-GDP | $2.59 \times 10^{-4}$ |
| SASA V9 SD | $4.20 \times 10^{-4}$ |
| SASA M72 SD | $4.75 \times 10^{-4}$ |
| RMSF I100 | $5.61 \times 10^{-4}$ |
| RMSD backbone-GDP | $1.37 \times 10^{-3}$ |
| Entropy quasi-harmonic | $1.47 \times 10^{-3}$ |
| SASA G60 AV | $1.99 \times 10^{-3}$ |
| SASA D69 SD | $2.61 \times 10^{-3}$ |
| SASA protein | $3.06 \times 10^{-3}$ |
| RMSF Q61 | $3.16 \times 10^{-3}$ |
| Coulomb 1,4 | $3.64 \times 10^{-3}$ |
| RMSF D69 | $3.72 \times 10^{-3}$ |
| SASA C12 AV | $4.45 \times 10^{-3}$ |
| RMSF G10 | $7.66 \times 10^{-3}$ |
| MSD bs*-backbone | $1.01 \times 10^{-2}$ |
| RMSF E63 | $1.19 \times 10^{-2}$ |
| SASA E63 AV | $1.77 \times 10^{-2}$ |
| SASA Q61 SD | $1.94 \times 10^{-2}$ |
| SASA H95 SD | $2.22 \times 10^{-2}$ |
| Rg protein | $2.69 \times 10^{-2}$ |
| SASA D69 AV | $4.65 \times 10^{-2}$ |
| SASA T35 SD | $4.84 \times 10^{-2}$ |

\*Abbreviations: AV, per average; bs, switch II binding site.

Another statistically significant feature associated with structural displacement was the mean square displacement (MSD) of the switch II binding site backbone (**Fig 2E**, **Table 1**). Like the protein–GDP complex, the MSD distribution for the switch II binding site backbone was shifted toward higher values in the treatment-sensitive systems (median: 4.82 vs 4.47 nm²) and showed greater variability (mean 4.73 nm² ± 0.86 nm² vs. 4.39 nm² ± 0.77 nm²), compared with the treatment-resistant group. This indicates that, on average, backbone atoms in the switch II binding region moved farther from their reference positions in the treatment-sensitive systems. In contrast, the radius of gyration (Rg), a measure of a protein compactness, showed a slight increase in the treatment-resistant group values compared to the treatment-sensitive systems (median: 1.57 with max Rg: 1.61 nm vs 1.56 and 1.59 nm, **Fig 2F**). Although the magnitude of this difference was small, it yielded a *P* value marginally below .05, suggesting borderline statistical significance.

Unlike global RMSD, which captures overall structural deviation, root mean square fluctuation (RMSF) provides insight into residue-level flexibility. **Figs 3A-B** shows RMSF values across the entire simulation time, with arrows marking peaks above 0.4 nm to highlight the most flexible residues. In both groups, residues T35, E63, and Y64 consistently exhibited the highest peaks. Additionally, P34 and E62 showed increased fluctuations in the treatment-sensitive systems, while E37 and Q61 showed higher flexibility in the treatment-resistant group. Among these, only Q61, E63, and Y64 exhibited statistically significant RMSF differences between groups (**Table 1**). Analysis of the most populated conformations revealed additional switch II binding site residues with significant group-wise RMSF differences, including M72, Q99, H95, I100, D69, and G10 (**Fig 3C**). For 8 out of the 9 reported residues, the treatment-resistant group had higher median RMSF values than the treatment-sensitive group, indicating greater structural flexibility.

**Fig 3.**
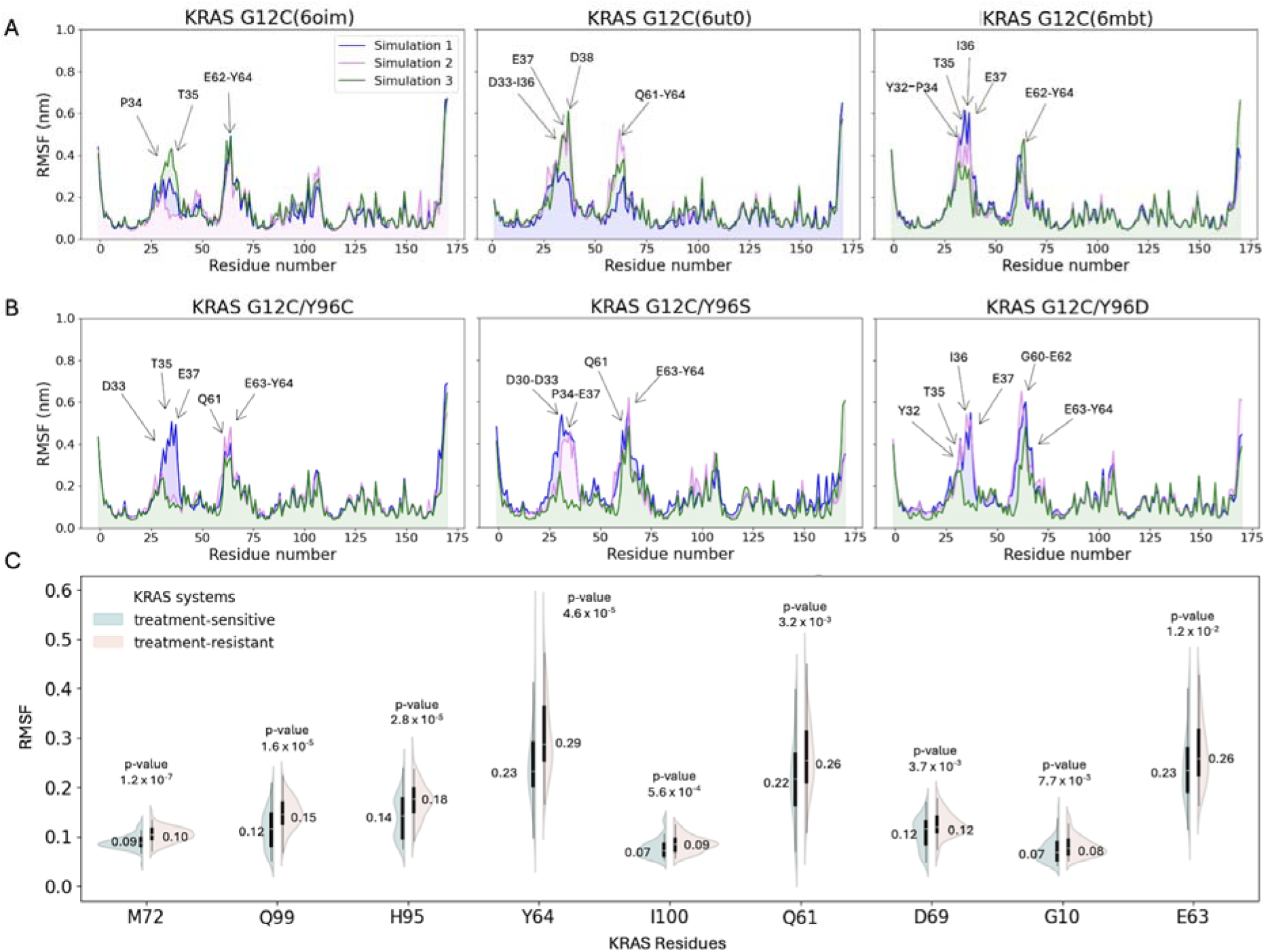
RMSF over the molecular dynamics simulation. Residues with RMSF >0.4 nm are indicated with arrows. (**A**) KRAS treatment-sensitive systems. (**B**) KRAS treatment-resistant systems. (**C**) RMSF data distribution for the individual residues in the switch II binding site ranked by the lowest *P* value, with annotated median values. Abbreviations: KRAS, Kirsten rat sarcoma viral oncogene homolog; RMSF, root mean square deviation.

The final structural feature analyzed was SASA. Regarding the total SASA of the protein in the most populated conformations, treatment-resistant systems exhibited slightly higher values (median: 97.45 nm²) compared with treatment-sensitive systems (95.95 nm²), indicating increased solvent exposure, statistically significant between groups (**Table 1**). To gain residue-level insight, SASA was further analyzed within the switch II binding site. Violin plots in **Figs 4A-B** highlight the top 5 residues with the lowest *P* values for both average (AV) and SD of SASA. Across these residues, median SASA values were consistently higher in the treatment-resistant group, suggesting increased solvent exposure. The most significant difference was observed in the SASA (SD) for residue 10 (*P* = 1.5 × 10⁻¹⁷), followed by the SASA (AV) of residue 95 (*P* = 1.8 × 10⁻¹²) and the SASA (SD) for residue 62 (*P* = 6 × 10⁻⁸), highlighting substantial variation in solvent accessibility between groups at the individual residue level.

**Fig 4.**
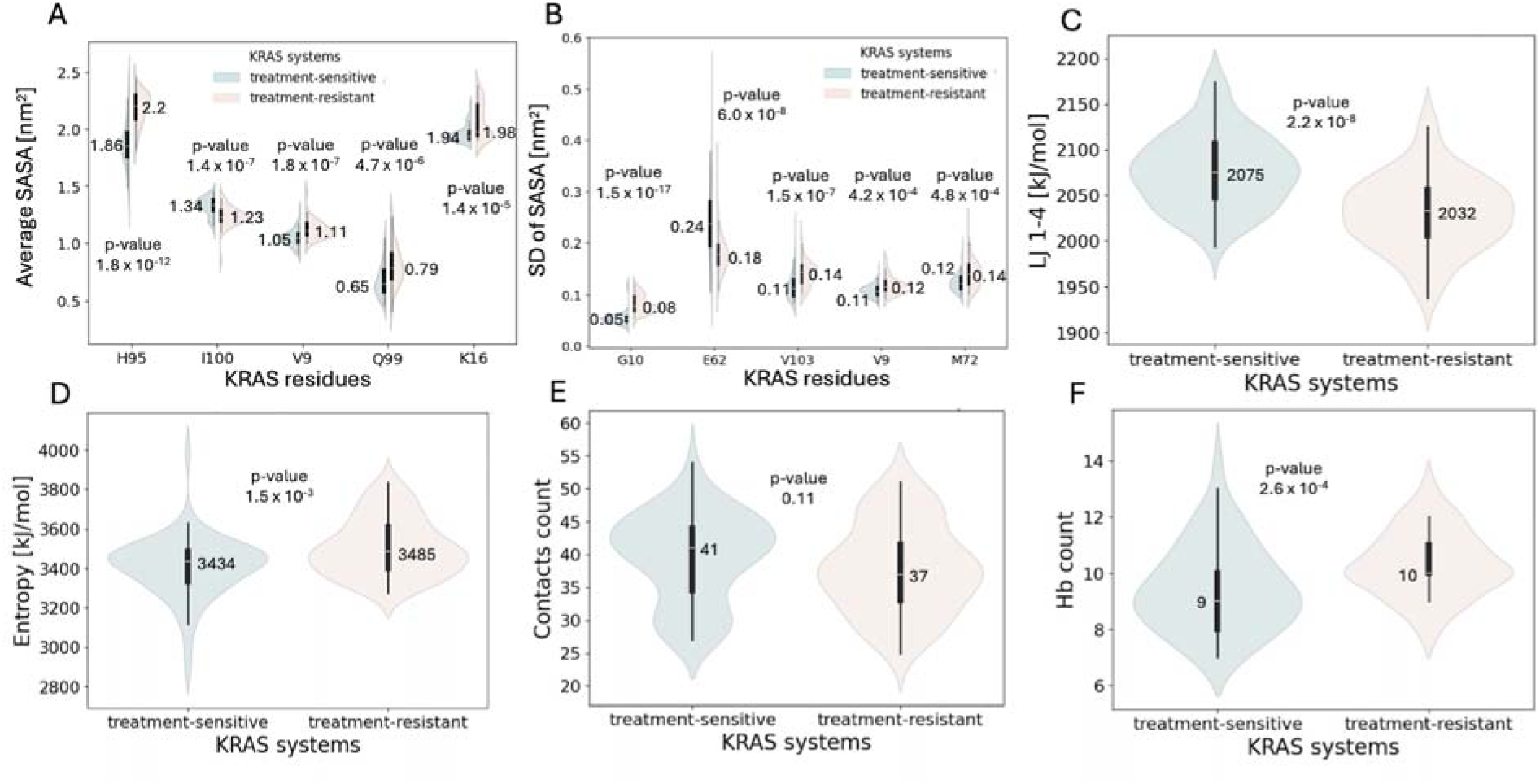
Data distribution of structural features, thermodynamic features and intra- and intermolecular contacts with annotated median and P values. (**A**) SASA (AV) of the switch II binding site residues ranked by the lowest P value. (**B**) SASA (SD) for the switch II binding site residues ranked by the lowest P value. (**C**) LJ 1,4 energy. (**D)** Entropy calculated according to the quasiharmonic equation. (**E**) Intramolecular contacts count between C12 and the rest of protein residues. (**F**) Hydrogen bonds count between the protein residues and GDP. Abbreviations: GDP, guanosine diphosphate; LJ, Lennard-Jones; SASA, solvent-accessible surface area.

The next group of features included energetic properties: potential energy, kinetic energy, and both short-range and 1,4 Coulomb and Lennard-Jones (LJ) interaction energies. Among these, LJ 1,4 energy and Coulomb 1,4 energy showed statistically significant differences between treatment-sensitive and treatment-resistant groups (**Table 1**). Lennard-Jones 1,4 energy had the third lowest *P* value among all features, with consistently higher values in the treatment-sensitive group (median: 2075 mean: 2077 kJ/mol) compared with the treatment-resistant group (2032 and 2031 kJ/mol), highlighting the difference in nonbonded interactions energies between the groups (**Fig 4C**). Conversely, Coulomb 1,4 energy was significantly elevated in the treatment-resistant systems (median: 23 369 vs 23 234 kJ/mol, data not shown).

To gain insight into thermodynamic properties of the systems, enthalpy and entropy were analyzed. Entropy calculated using the quasiharmonic approximation showed a statistically significant difference between treatment-sensitive and treatment-resistant groups (**Table 1**, **Fig 4D**) with elevated values in the treatment-resistant systems (median: 3485 vs 3434 and mean: 3507 vs 3407 kJ/mol). This finding aligns with the generally increased conformational flexibility in the treatment-resistant group, as indicated by structural features such as RMSF and SASA.

Intra- and intermolecular contacts were also analyzed. Intramolecular interactions included contacts between the protein and individual amino acid residues, as well as hydrogen bond formation within the protein. Intermolecular interactions focused on hydrogen bonds formed between the protein and GDP. The first group of contacts included residue 12, as cysteine at this position serves as the covalent binding partner for small-molecule inhibitors targeting the KRAS G12C mutant. A higher number of contacts between C12 and the rest of the protein were observed in the treatment-sensitive systems, compared with the treatment-resistant ones (median: 41 vs 37, **Fig 4E**); however, this difference was not statistically significant. Among hydrogen bond–related features, the number of protein-GDP hydrogen bonds differed significantly between the 2 groups (**Table 1**). The treatment-resistant systems exhibited a higher median hydrogen bond count (median:10, range 8-13) than the treatment-sensitive group (9 and range 7-14), indicating more frequent protein-GDP interactions in the treatment-resistant group on average, although the treatment-sensitive group exhibited a more disperse range (**Fig 4F**). This finding is correlated to more stable GDP binding pocket in treatment-resistant group (**Fig 2D**), which allows for more stable occurrence of hydrogen bonds on average. It was reported in previous studies [30] that both stronger and weaker GDP binding is possible for KRAS secondary mutants.

Statistical analysis suggested that the relationships between molecular features are complex, with no single trend consistently distinguishing the 2 groups. For instance, the treatment-resistant systems showed increased flexibility in the switch II binding site region, as indicated by RMSF and SASA, while the treatment-sensitive systems exhibited greater mobility in the same region’s backbone, as measured by MSD. These complex findings highlight the structural variability at the residue level, making it challenging to draw definitive conclusions about the overall direction or significance of individual features.

### Machine-learning models

Given the complexity of MD simulations data, we applied ML to uncover multivariate patterns that may not be apparent through univariate analysis alone, and to identify the most informative features capable of distinguishing treatment-resistant mutants from treatment-sensitive ones (**Fig 5A**). All predictive ML models (LR, RF, and SVM), achieved high classification performance, with average accuracy and area under the curve exceeding 90% (**Table 2**, **Fig 5B**). Model stability across methods was confirmed through 10-fold cross-validation. Hyperparameter tuning using GridSearchCV yielded parameter values close to the defaults (**Table 2**), further indicating robust and generalizable performance.

**Fig 5.**
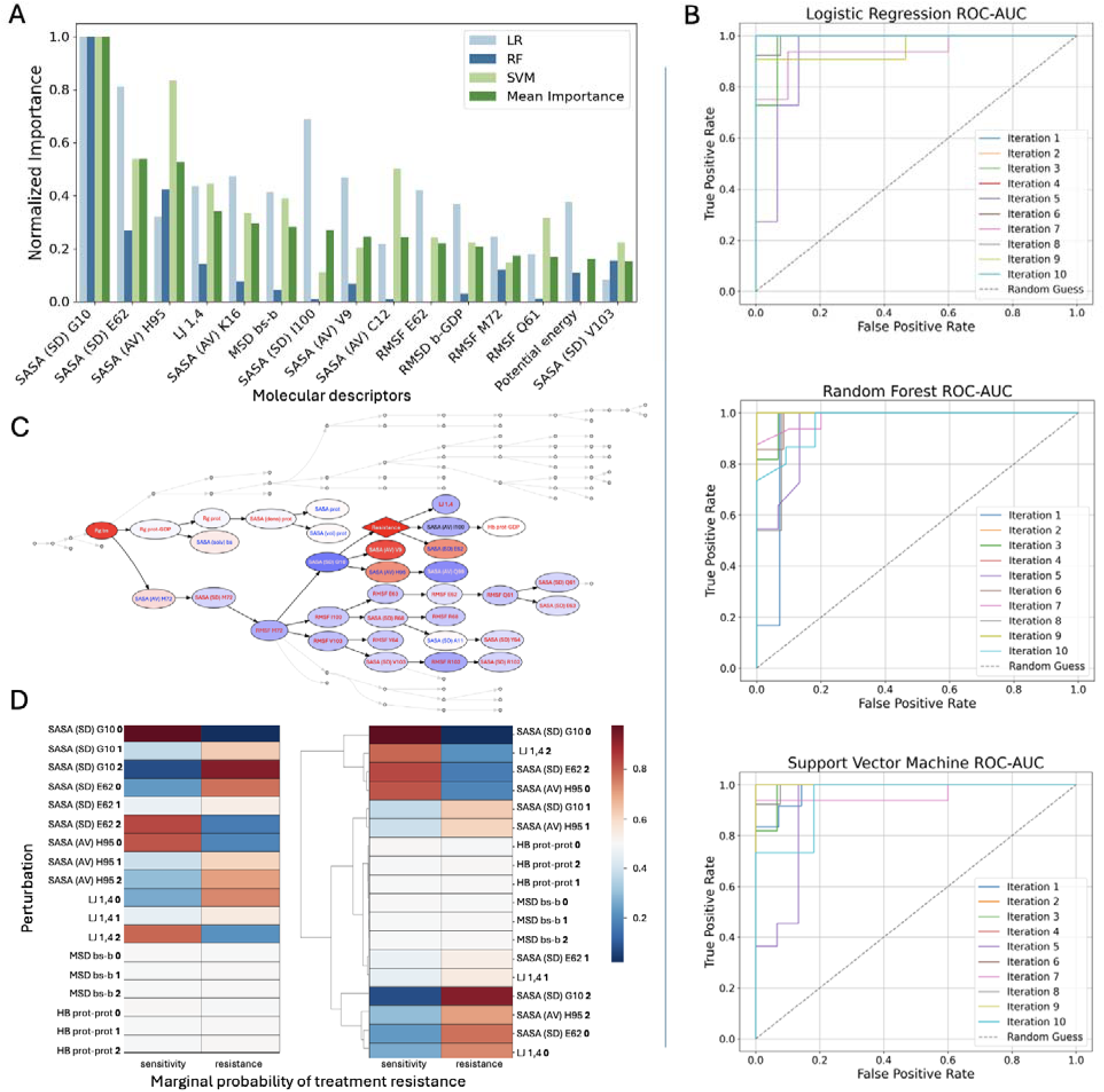
Machine-learning analyses. (**A**) The top important features selected by the machine learning models: LR, RF, and SVM, ranked in the descending order by mean importance, normalized on a 0 to 1 scale. (**B**) Performance of the ML models. ROC AUC curves for LR, RF, and SVM models. (**C**) BN: a subset of the network highlighting the top features is shown. Node colors correspond to the state of each feature when the *resistance* node is fixed to state 1, with red indicating higher marginal value and blue indicating lower marginal value. (**D**) Comparison of how different feature-state combinations influence the likelihood of treatment resistance outcome in BN. Each cell shows the marginal probability of treatment resistance when a specific molecular descriptor is fixed at a given state (0, low; 1, medium; and 2, high). Abbreviations: AUC, area under the curve; BN, Bayesian network; LR, logistic regression; ML, machine learning; RF, random forest; ROC, receiver-operating characteristic; SVM, support vector machine.

**Table 2.**
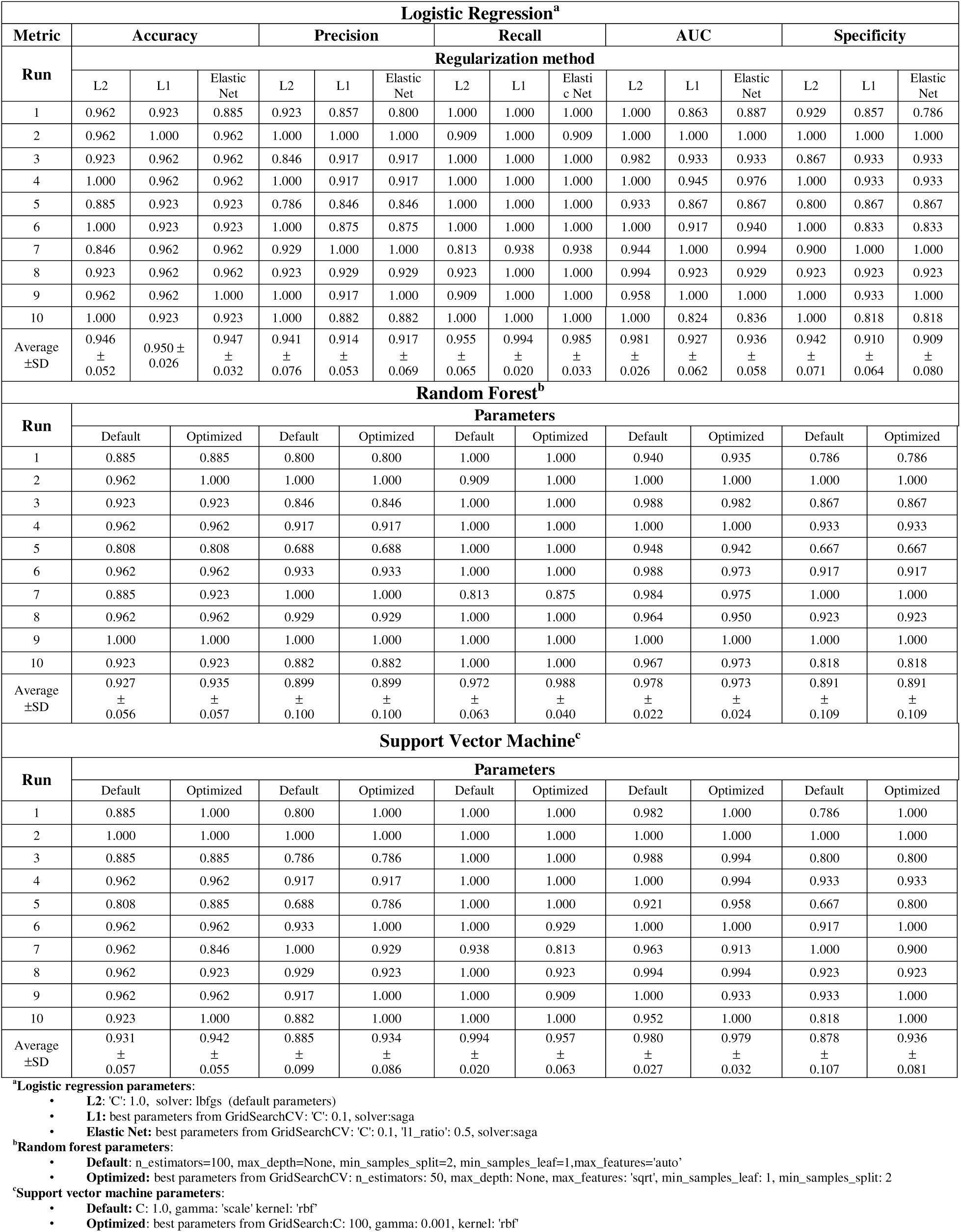
Performance of the ML Models. Logistic Regression (LR), Random Forest (RF) and Support Vector Machine (SVM) in terms of standard metrics: accuracy, precision, recall, area under the curve and specificity. Evaluated using 10-fold cross-validation.

Across all ML models, the SASA (SD) for residue G10 (**Fig 5A**) was consistently ranked as the most predictive feature differentiating treatment-resistant from treatment-sensitive KRAS systems. This result aligns with the Mann–Whitney U test, where this feature had the lowest *P* value among all descriptors, indicating the strongest statistical significance. The SASA (SD) for residue G10, was elevated in treatment-resistant systems (**Fig 4B**), along with its RMSF, which was also statistically significant (**Table 1**).

Other top-ranked features included the SASA (AV) for H95 and the SASA (SD) for E62, both of which also had among the lowest *P* values. The SASA (AV) for H95 was notably higher in treatment-resistant systems, similar to the residue G10 (**Fig 4A**). In contrast, the SASA (SD) for E62 was elevated in treatment-sensitive systems (**Fig 4B**). Additionally, its RMSF showed high discriminatory power in the LR model, further highlighting its contribution to conformational dynamics relevant to treatment response.

Additional features prioritized by the models included LJ 1,4 energy, the SASA (AV) of K16, and the MSD of the switch II binding site backbone. All were statistically significant, with LJ 1,4 energy ranked as the third most significant feature overall. LJ 1,4 energy values and MSD of the switch II binding site backbone were both more concentrated at higher levels in the treatment-sensitive systems (**Figs 4C and 2E**, respectively), whereas the SASA (AV) of K16 was elevated in the treatment-resistant systems (**Fig 4A**). For validation, we ran two additional ML models (XGBoost and PLS-DA), which remained in agreement with the results. Models’ performance and evaluation can be found in the Supplementary Material (**S1 and S2 Tables**, **S1 and S2 Figs**).

Bayesian Network models helped to understand the underlying structure of relationships between the molecular features. First, BN inference was used to explore how treatment response status influences the probabilistic behavior of molecular descriptors. When the treatment resistance node was fixed to either the sensitive or resistant state, the resulting shifts in marginal probabilities revealed that treatment resistance exerts a structured, nonrandom influence on specific network regions. The most strongly affected descriptors included the SASA (SD) for G10 and E62, the SASA (AV) of H95, and LJ 1,4 energy (**Fig 5C**), aligning with the features previously identified by ML models.

To assess whether these descriptors could, in turn, influence treatment response, each was fixed to one of its discretized states, and the resulting marginal probabilities of the treatment resistance node were computed. For comparison, 2 additional descriptors, the MSD of the binding site backbone and the number of intraprotein hydrogen bonds, were also tested because they showed little change during resistance-driven inference. This analysis revealed several nonlinear, descriptor-specific effects. Specifically, reduced values of SASA for G10 (SD) and H95 (AV), and increased values of SASA for E62 (SD) and LJ 1,4 energy, were associated with higher marginal probabilities of treatment sensitivity (**Fig 5D**). Conversely, the opposite descriptor states consistently shifted probabilities toward treatment resistance, reinforcing their importance in shaping the resistance phenotype (**Fig 5D**). These findings are consistent with the ML results regarding the molecular features distinguishing between the groups.

To visualize the features identified by ML models, the most populated MD conformations from both treatment-sensitive and treatment-resistant KRAS systems were examined regarding the residues G10, E62, and H95 conformations (**Fig 6**). Residues E62 and H95 exhibited distinct conformational dynamics across the most populated clusters, which may explain the differences in solvent exposure values and suggest potential alterations in ligand accessibility that warrant further investigation. Residue G10 was located within a highly flexible region, where neighboring residues likely contribute to the dynamic variability of its solvent exposure.

**Fig 6.**
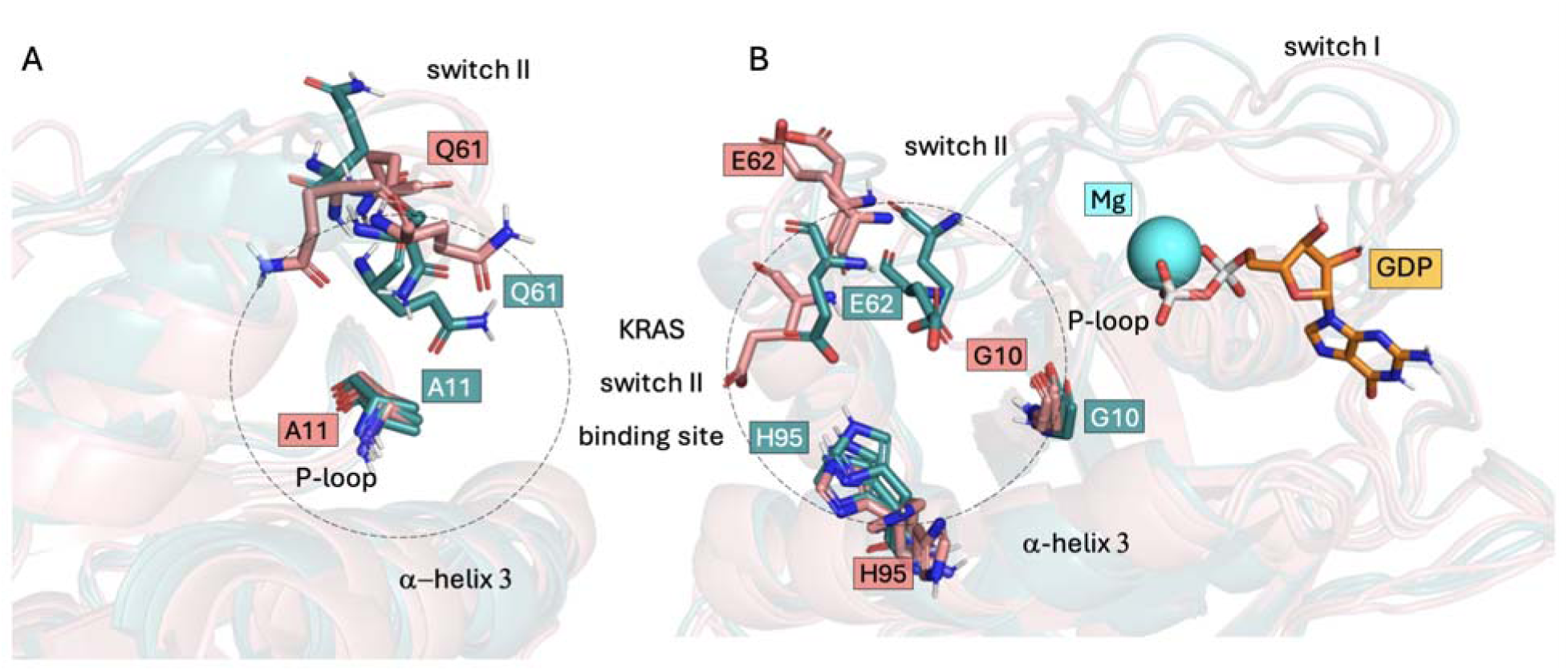
Switch II binding-site dynamics. Aligned most populated conformations from MD simulations with important residues shown in sticks. (**A**) Side view. Residue A11 was removed from the ML predictions because of high correlation with target variable and Q61 is known for its catalytic activity. (**B**) Front view. The most important residues indicated by ML models (G10, E62, and H95). Treatment-sensitive systems (KRAS G12C) in teal, treatment-resistant systems (KRAS G12C/Y96C, KRAS G12C/Y96D, and KRAS G12C/Y96S) in salmon. Three examples from each class are shown for clarity. Abbreviations: KRAS, Kirsten rat sarcoma viral oncogene homolog; MD; ML, machine learning.

Overall, univariate statistical analysis and ML models (LR, RF, SVM, and BN) identified a consistent set of molecular features that differed significantly between treatment-sensitive and treatment-resistant systems. There was strong agreement across methods regarding feature relevance, particularly for structural descriptors. The most influential features were predominantly related to SASA within the switch II binding site. Across all ML models, the SASA (SD) for G10 emerged as the top discriminatory feature, followed by the SASA (AV) of H95, the SASA (SD) for E62, and LJ 1,4 energy. These findings highlight the central role of residue-level flexibility, solvent exposure and nonbonded interactions in KRAS mutation-induced drug resistance.

## Discussion

In this study, the unit of biological variation was the KRAS variant; we analyzed four variants (G12C; G12C/Y96C, G12C/Y96S, G12C/Y96D) using 126 conformers from multiple PDB starts, replicas, and RMSD clusters, intentionally focusing on, although limited, yet clinically reported, resistance-associated double mutants within the inhibitor-binding site to provide hypothesis-generating results that we will expand with additional sensitive variants in future work. It is important to underline that all MD were performed in the unbound (apo) state; no complexes with inhibitors were simulated. Thus, any conclusions are restricted to conformational/topological shifts and do not establish inhibitor-specific resistance mechanisms. These unbound-state results motivate future testing with representative inhibitors but do not by themselves define resistance.

Our approach identified several residues (G10, K16, E62, and H95) as important contributors to treatment resistance in our models, consistent with their known roles in inhibitor recognition. For example, it has been shown that adagrasib forms a salt bridge with E62 and hydrogen bonds with H95 and K16, while its cyanomethyl group interacts with G10, displacing a conserved water molecule [17]. While sotorasib also forms hydrogen bonds with K16, it exploits a specific H95 orientation enhancing binding affinity to KRAS G12C, forms hydrogen bonds E63, and π- π interactions with Y96 [16]. Additionally, G10 solvation has been extensively discussed in previous structural and MD studies [17, 30, 41]. In our apo-state simulations, resistant systems showed increased solvent exposure of G10 and H95 residues and reduced flexibility of E62, suggesting altered solvation patterns and local rigidity around the binding site. These conformational features, even in the absence of ligand, pre-organize resistant mutants into states less compatible with optimal inhibitor binding.

Interestingly, while our ligand-free simulations of secondary resistant KRAS variants (e.g., G12C/Y96C/D/S) revealed mutation-dependent changes within the local contact network around G10, K16, E62, and H95, independent ligand-bound MD studies have reported parallel shifts in the contact frequencies of adagrasib and sotorasib with these same residues in mutants such as G12C/R68S, G12C/K16T, G12C/Y96C, and G12C/Y96D [30], together supporting a model in which resistance stems from subtle, mutation-induced conformational changes that pre-organize the inhibitor-binding interface in the apo state and subsequently alter drug contacts upon binding.

In summary, we developed a framework that integrates computational chemistry methods with machine learning algorithms, using KRAS mutation-induced resistance as a case study. Within this framework, a dataset of comprehensive molecular descriptors, capturing structural, energetic, thermodynamic, and contacts-based properties was derived from MD simulations and analyzed by multiple methods. Predictive ML models, LR, RF, and SVM classified treatment response based on molecular descriptors, while the BN revealed probabilistic dependencies between features, enhancing interpretability. Together, these approaches identified subtle yet meaningful differences in the dynamics of KRAS treatment-sensitive and treatment-resistant systems, particularly in residue-level flexibility and solvent exposure. The conformational behavior of key residues within the switch II binding site, as demonstrated in our results, could be considered in the design of inhibitors for KRAS secondary mutants, resistant to sotorasib and adagrasib, potentially improving therapeutic outcomes. Our study highlights the potential of ML to address the complexity of mutation-induced drug resistance.

To date, the integration of protein flexibility or MD-derived features into ML models has not been applied to problems as complex as mutation-induced drug resistance. Our approach captures mutation-driven intrinsic conformational changes, not ligand-specific effects. This reduces mechanism specificity but improves generalizability to other phenotypes and targets. By incorporating protein dynamics before ligand-based analyses, it becomes possible to uncover unique, targetable features that support the development of more selective and effective inhibitors. Given the widespread impact of mutation-induced resistance across cancer types, we propose extending this framework to other clinically relevant resistance-associated targets in the future.

## Methods

### Computational chemistry

To investigate the dynamic behavior of KRAS mutants, we conducted all-atom MD simulations of the inactive KRAS conformations bound to Mg^2+^ and GDP. The initial protein structures were obtained from the Protein Data Bank (PDB) [42] under accession codes: 6OIM [17], 6UT0 [16], and 6MBT [43], and prepared before MD simulations. For the KRAS G12C mutant bound to the covalent inhibitor sotorasib (PDB ID: 6OIM), crystallographic water molecules were removed together with the ligand. Missing amino acid residues 105-107 were reconstructed using MODELLER 10.5 [44]. Three engineered mutations, C51S, C80L, and C118S [15], present in the structure were reverted back to cysteines using the PyMOL 3.0 mutagenesis panel [40]. The protein was capped with an acetyl group at the N-terminus and with a methyl amide at the C-terminus using PyMOL’s builder module [40]. This structure served as the basis for building KRAS double mutant models, which have not yet been experimentally resolved. Subsequently, KRAS G12C (PDB ID: 6OIM) was manually mutated in PyMOL to construct models of the double mutants G12C/Y96C, G12C/Y96S, and G12C/Y96D. For the wild-type (WT) KRAS structure (PDB ID: 6MBT) and the KRAS G12C mutant bound to the covalent inhibitor adagrasib (PDB ID: 6UT0), the same preparation steps were applied. Additionally, the WT KRAS structure was mutated at position 12, from glycine to cysteine, to model an additional KRAS single-mutant system.

All protein models were processed using the CHARMM36 force field [45] in GROMACS [46]. Hydrogen atoms were added to each protein structure, and the protonation states of the residues were assigned at neutral pH. The GDP molecule was parametrized separately with the CHARMM General Force Field [47]. To prepare the systems for energy minimization, the complexes were placed in cubic boxes, extending at least 10 Å from the protein. The solvent was modeled using the CHARMM-modified TIP3P water model [45]. The protein charges were neutralized by adding the appropriate amount of Na+ ions, and physiological NaCl salt concentration 0.1M was ensured. Energy minimization was performed using the steepest descent algorithm [48].

Next, NVT equilibration was conducted for 100 ps to stabilize the system’s temperature at 310 K with position restraints on protein and ligand heavy atoms to allow the solvent and ions to relax. Temperature was maintained at 310 K using the V-rescale thermostat [49] with separate coupling groups for protein and GDP ligand, and solvent/ions. H-bonds were constrained using LINCS [50], enabling the 2 fs timestep. This was followed by 250 ps of NPT equilibration, continuing from the NVT step, with the same position restraints and temperature control, while pressure was maintained at 1 bar using isotropic Berendsen coupling [51]. Production MD were performed for 200 ns with a 2 fs timestep. Temperature was maintained at 310 K using the V-rescale thermostat, and pressure was maintained at 1 bar using isotropic Parrinello-Rahman coupling. Coordinates and energies were saved every 10 ps. Each system underwent 3 replica runs, with different random seeds set before each simulation run. The water shell confirming octahedral arrangement of water molecules around Mg^2+^ ion is shown in **S3 Fig**.

To extract representative conformations from the trajectories, we performed trajectory clustering on the 50–200 ns portion of each simulation using the GROMOS [52] method. This method counts neighboring structures within a specified cutoff and identifies the representative frame as the one with the smallest average root mean square deviation (RMSD) to all its neighbors. Clustering was performed for 6 different Cα RMSD cutoffs, ranging from 0.10 nm to 0.15 nm, to find reasonable distribution of frames for each system. Representative structures from the top 7 clusters (∼90% of the trajectory) were extracted from each of the 3 replicas per system for molecular descriptors calculations. This resulted in 126 conformations across all 6 systems. Specific elements of each system were treated separately in the calculations, including extended switch II binding site residues (residues 9, 10, 11, 12, 16, 34, 35, 37, 58, 59, 60, 61, 62, 63, 64, 68, 69, 72, 95, 96, 99, 100, 102, and 103), which were defined as residues within 5Å of any site point identified in the KRAS G12C switch II binding site by the SiteMap module [53, 54] of Schrödinger Suite and included motifs from P-loop, switch I region, and adjacent helices. A full list of molecular descriptors, grouped into categories, is provided in **Table 3**.

**Table 3.** The complete list of the molecular descriptors extracted from MD simulations.

| **Molecular dynamics simulation descriptors** |  |
| --- | --- |
| **Calculation type** | **System elements** |
| **Structural descriptors** |  |
| Root Mean Square Deviation (RMSD, nm)* | Calculated as a function of time for:<br>protein backbone<br>switch II binding site residues (backbone and sidechains - heavy atoms)<br>GDP and protein backbone<br>Mg<sup>2+</sup> ion and protein backbone |
| Radius of gyration (Rg, nm)* | Calculated as a function of time for (all-atoms):<br>protein<br>protein-GDP<br>switch II binding site residues |
| Root Mean Square Fluctuation (RMSF, nm) | Calculated per switch II binding site residue over the trajectory frames extracted for each cluster. |
| Solvent accessible surface area (SASA, nm<sup>2</sup>) using the Double Cubic Lattice Method[55] | Total solvent accessible surface area, estimated solvation free energy, and total volume and density was calculated as a function of time for (all-atoms):<br>protein<br>switch II binding site residues<br>Additionally, the average solvent accessible surface area and its standard deviation was calculated per each switch II binding site residue over the trajectory frames extracted from each cluster. |
| Mean Square Displacement (MSD, nm<sup>2</sup>) | Calculated as a function of time for:<br>protein backbone<br>switch II binding site residues (backbone)<br>switch II binding site residues (sidechains-heavy atoms). |
| **Energetic features** |  |
| Potential energy, total energy, kinetic energy, short-range and 1,4 Coulomb and Lennard-Jones (LJ) energies (kJ/mol) | Extracted for representative frame from the most populated conformational clusters. |
| **Thermodynamic features** |  |
| Entropy estimate based on the quasi-harmonic approach[56] and Schlitter's formula[57] (kJ/mol) | Estimated as an evolution of the system up to each cluster's representative timeframe. |
| Enthalpy (kJ/mol) | Extracted for representative frame from the most populated conformational clusters. |
| **Intra- and intermolecular contacts** |  |
| Hydrogen bonds count | Calculated as a function of time for (all atoms):<br>within protein<br>between protein and GDP<br>within switch II binding site residues |
| Number of contacts within 6 Å | Calculated as a function of time for (heavy atoms):<br>between residue 12 and protein<br>between residue 12 and switch II binding site residues |

\*First frame of the production run was used as the reference conformation.

### Statistical and ML methods

ML approach was implemented to identify the molecular descriptors distinguishing between treatment-sensitive and treatment-resistant KRAS mutants. The dataset consisted of 3 treatment-sensitive (resistance = 0) and 3 treatment-resistant (resistance = 1) KRAS systems, each contributing 63 samples, for a total of 126 samples described by 101 molecular features. To reduce redundancy, highly target-correlated and intercorrelated features (correlation >75%) were removed, along with all descriptors directly related to residue 96, the site of the secondary mutation, to avoid bias toward known changes. The remaining features were standardized using Z-score scaling to ensure comparable value ranges across inputs.

Before model training, we performed univariate statistical analysis. Feature distributions were tested for normality using the Shapiro–Wilk test [58], and the Mann–Whitney U test [59] was used to assess statistical significance between the feature data distribution in the treatment-sensitive and treatment-resistant groups. Violin plots [60, 61] were used to visualize feature distributions, with central boxes indicating the interquartile range (IQR, 25th–75th percentile), median lines, and whiskers extending 1.5× the IQR.

Since the relationships between features and resistance were not assumed to be strictly linear, 3 supervised ML models were applied: logistic regression (LR), random forest (RF), and support vector machine (SVM), covering both linear and nonlinear classification paradigms. Hyperparameter optimization was performed using GridSearchCV [62] with 5-fold cross-validation. For LR, the L1 (Lasso), L2 (Ridge), and Elastic Net [62, 63] regularizations were tested. The regularization strength C was varied (0.01, 0.1, 1, 10, 100) for all models, while Elastic Net additionally included tuning of the l1_ratio (0.1, 0.5, 0.7, 1.0). For the SVM classifier, the grid included C (0.01, 0.1, 1, 10, 100), kernel (linear, rbf, poly), gamma (0.001, 0.01, 0.1, 1), and degree (2, 3, 4). For the RF model, the parameters tested were n_estimators (50, 100, 150), max_depth (none, 10, 20), min_samples_split (2, 5, 10), min_samples_leaf (1, 2, 4), and max_features (sqrt, log2, none). Feature importance was determined by using absolute coefficients for LR, total Gini impurity reduction for RF, and permutation importance for SVM. Each model was trained on 80% of the data and tested on the remaining 20%, with performance evaluated using 10-fold cross-validation to ensure repeatability and prevent overfitting [63]. All analyses were conducted using Python 3.11.11[64].

To analyze probabilistic relationships among molecular descriptors, we used a Bayesian Network (BN), an interpretable ML model that represents joint probability distributions as a directed acyclic graph. A universal BN was constructed by combining treatment-sensitive and treatment-resistant KRAS datasets using BNOmics [65], with 100 random restarts to ensure convergence and structural robustness. Molecular descriptors were discretized into 3 categories, low (0), medium (1), and high (2), using a maximum entropy algorithm [65]. The resulting network captured both direct associations with treatment response and higher-order dependencies among features. Two types of inference analyses were performed using a belief propagation algorithm. First, to assess how treatment response affects molecular descriptors, the treatment resistance node was fixed to either sensitive (0) or resistant (1), and the resulting posterior marginal probabilities were computed for each feature. These distributions quantify how each node’s probability shifts in response to the fixed phenotype. Second, to evaluate how selected molecular descriptors influence treatment response, each was fixed to one of its discretized states (0, 1, or 2), and the resulting marginal probability of the treatment resistance node was inferred. These probabilities were compiled into a heatmap to visualize the effect of each perturbation.

## Data and code availability

The GitHub repository containing the code and data can be found at: https://github.com/Molecular-AI-Group/KatMizgalska/tree/KRAS-MD-descriptors

## Author contributions

K.M. performed molecular dynamics simulations, conducted univariate statistical analyses and predictive ML modeling and wrote and edited the manuscript. K.U. and S.B. performed and described all analyses related to the BN. A.K designed the study in consultation with E.H., D.I. and W.G., supervised it, and provided guidance on result interpretation and manuscript editing. All authors reviewed and approved the final manuscript.

## Acknowledgments

Molecular dynamics simulations were performed using HPC computing cluster at H. Lee Moffitt Cancer Center and Research Institute. Editorial assistance was provided by Moffitt Cancer Center’s Office of Scientific Publishing by Gerard Hebert.

## Supporting Information

**S1 Fig.**
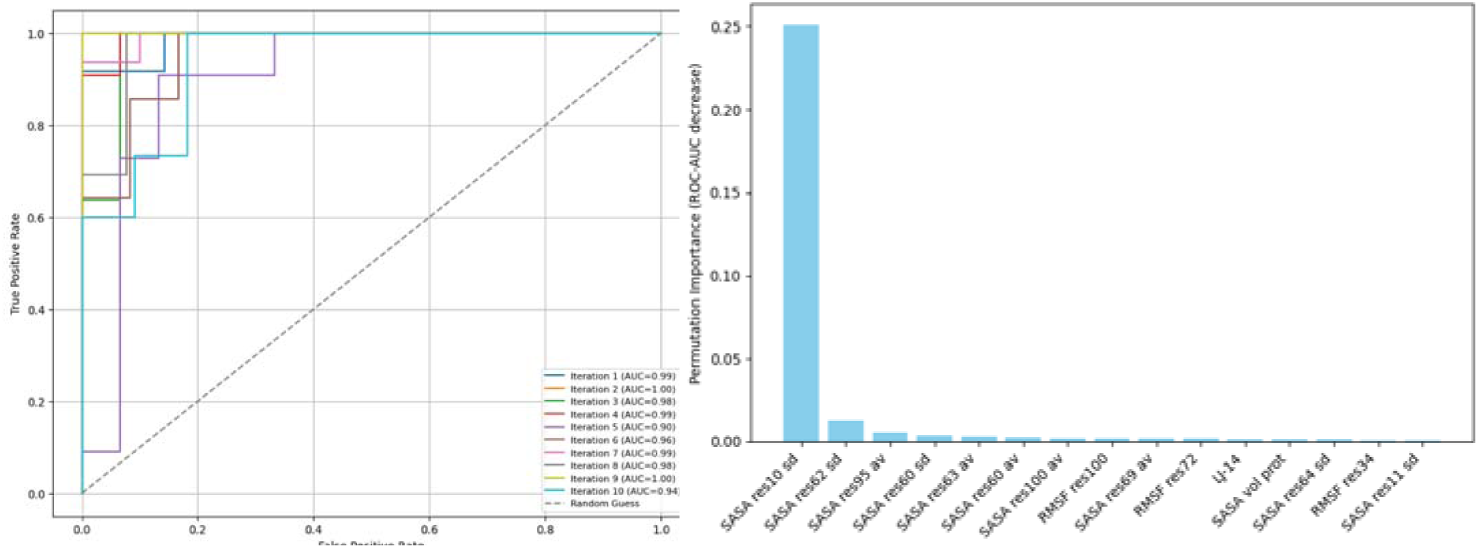
XGBoost model performance. **Left**: XGBoost ROC-AUC across 10 iterations. **Right**: top 15 Feature Importance averaged over 10 iterations calculated using a Permutation technique. This model, even after small-data set adjustment, still relies on one dominant feature SASA (SD) of G10. It agrees with the models studied in the publication.

**S2 Fig.**
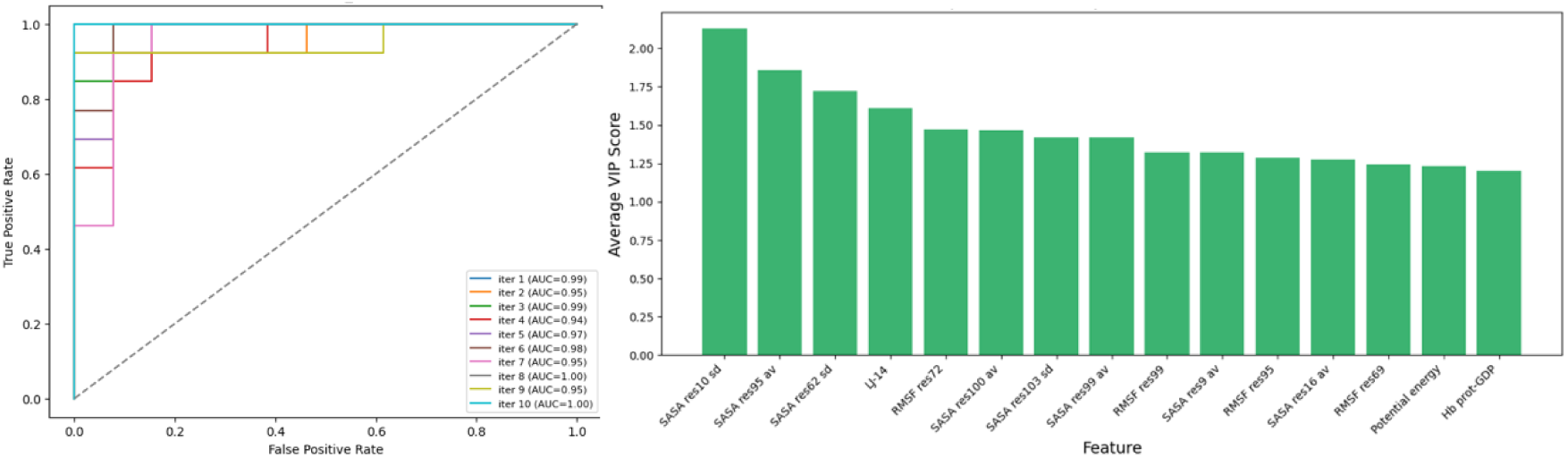
PLS-DA model performance. **Left**: PLS-DA ROC-AUC across 10 iterations. **Right**: top 15 Feature Importance averaged over 10 iterations calculated using a Variable Importance in Projection (VIP) scores, which estimate each feature contribution to the production. This model does not rely heavily on a few features, but consensus between several equally important. The result agrees with the models studied in the publication.

**S3 Fig.**
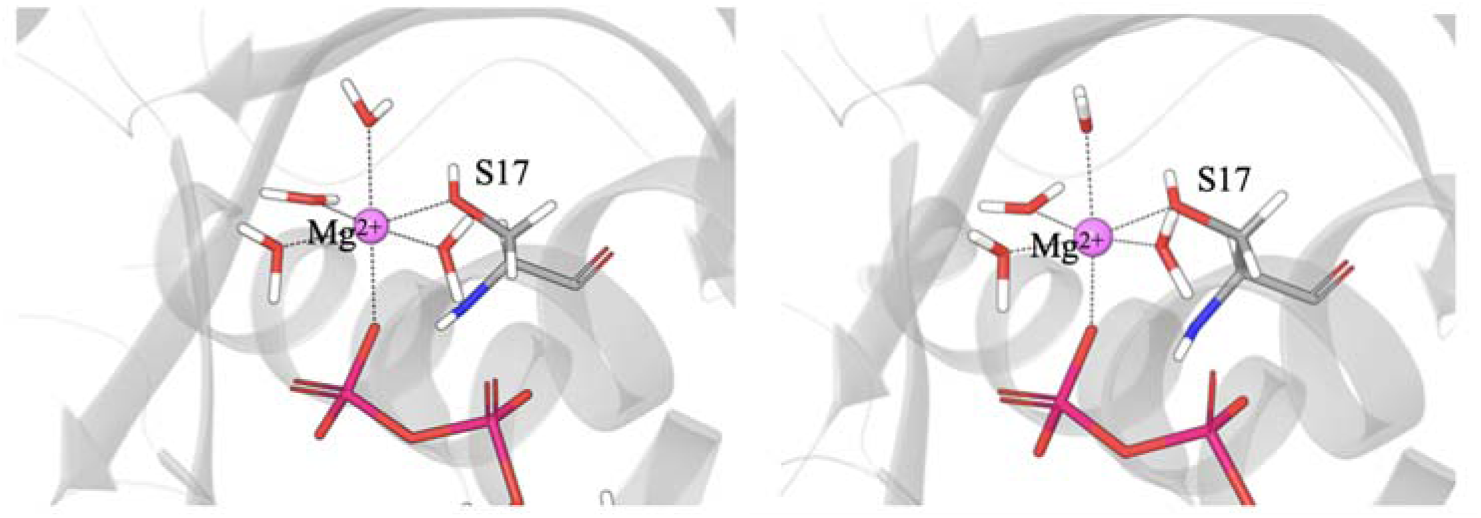
Water shell around Mg^2+^. **Left**: initial structure (PDB ID: 6OIM, X-ray). **Right:** system’s starting structure after the NPT equilibration ready for the simulation run.

**S1 Table.**
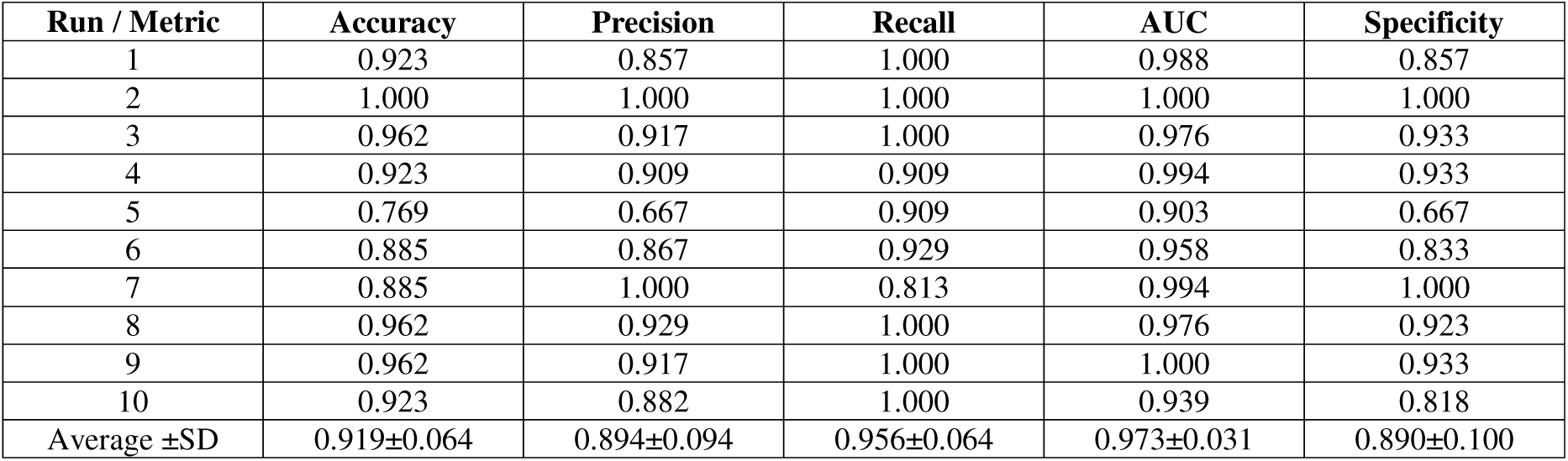
XGBoost model evaluation.

**S2 Table.**
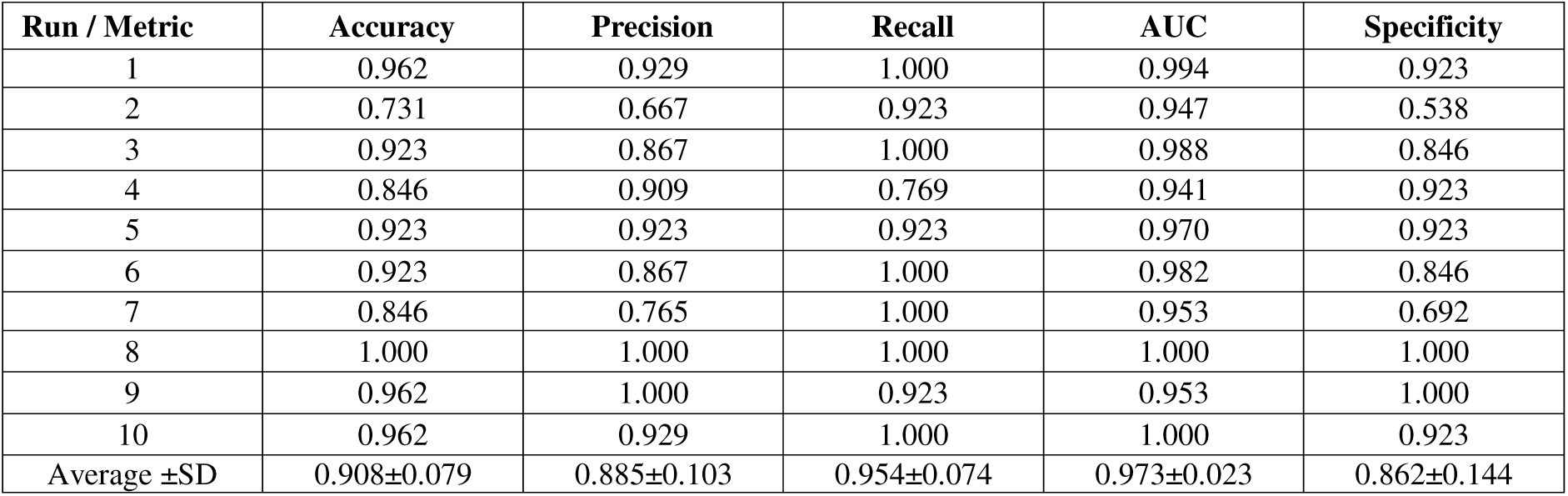
PLS-DA model evaluation.

